# Estimation of the instantaneous spike train variability

**DOI:** 10.1101/2023.11.19.567509

**Authors:** K. Rajdl, L. Kostal

## Abstract

The variability of neuronal spike trains is usually measured by the Fano factor or the coefficient of variation of interspike intervals, but their estimation is problematic, especially with limited amount of data. In this paper we show that it is in fact possible to estimate a quantity equivalent to the Fano factor and the squared coefficient of variation based on the intervals from only one specific (random) time. This leads to two very simple but precise Fano factor estimators, that can be interpreted as estimators of instantaneous variability. We derive their properties, evaluate their accuracy in various situations and show that they are often more accurate than the standard estimators. The presented estimators are particularly suitable for the case where variability changes rapidly.

## 1. Introduction

In the nervous system, information is transmitted by sequences of action potentials (spike trains). However, it is still not completely clear how exactly information is encoded. It is generally accepted that the timing of the firing of spikes is important, not their shape or size (Gerstner & Kistler, 2002; Rieke *et al*., 1999). Moreover, most information is apparently encoded by the spike rate, defined, for example, as the average spike count in a time window (Adrian & Zotterman, 1926; Perkel & Bullock, 1968). This concept is called rate coding. However, the idea that information is encoded by rate characteristics alone is simplified, different spike trains may have the same rate but carry different information based on the specific timing of the spikes (Shadlen & Newsome, 1994; Stein *et al*., 2005; Christodoulou & Cleanthous, 2011). Codes that also assume some rate-independent behavior of the spike trains are called temporal codes. The true coding method may be very complex, but it is reasonable to increase the complexity of the studied codes in steps, and the first step above rate coding is probably variability coding (Perkel & Bullock, 1968).

In most situations, the term variability is used to describe how distant some values, such as interspike intervals (ISIs) or spike counts, are from each other or from a typical value (Kostal *et al*., 2013). It is often measured by quantities based on statistical variance. Note that variability is a different concept from randomness. These terms may seem similar, but randomness focuses on how well some values are predictable and is mostly measured by entropy (Kostal *et al*., 2007; Rajdl *et al*., 2017).

The two most commonly used characteristics for measuring spike train variability are the coefficient of variation (CV), based on the variance of the lengths of the ISIs, and the Fano factor (FF), based on the variance of the number of spikes in a time window (Ditlevsen & Lansky, 2011; Stevenson, 2016; Rajdl *et al*., 2020; D’Onofrio *et al*., 2019). They are used, e. g., to measure the variability of real neuronal data (Festa *et al*., 2021; de Ruyter van Steveninck *et al*., 1997) or to study the neuronal variability using theoretical neuronal models (Protachevicz *et al*., 2022; Christodoulou & Bugmann, 2001). Note that these characteristics are used not only for neuronal data but also to measure the variability of data corresponding to point processes (sequences of times of some events) in many other fields (Yuan *et al*., 2012; Telesca *et al*., 2007; Contreras-Uribe *et al*., 2017). Their standard estimators have been well studied, and it is known that both have problems with bias in estimation from limited data. Specifically, the estimator of CV has a bias depending on the number of ISI observations (it is only asymptotically unbiased), and moreover, the observed ISIs are often right-censored (as they are limited by a fixed time window), which introduces another type of bias (Nawrot *et al*., 2008). The length of the observation window also strongly influences the estimator of FF, since FF by definition depends on the length of the observation window. The goal is often to estimate the limit of FF for a window length that goes to infinity. However, as in real experiments the window is always finite, this also leads to a bias (Rajdl & Lansky, 2014; Rajdl *et al*., 2020). The standard estimators of CV and FF are therefore not very suitable for estimating short-term (instantaneous) variability. An important fact is that for spike trains modeled as renewal processes it holds FF = CV^2^, assuming an infinite observation time window (Nawrot, 2010). This leads to seeing FF and CV^2^ as two ways to measure (estimate) the same quantity and increases the importance of their common value FF. In this paper we mostly write about the estimation of FF, but it could certainly be seen also as the estimation of CV^2^. Even in other papers these two terms are used interchangeably (Shuai *et al*., 2002). Note that there are other useful measures of variability. For example, CV2 was proposed in (Holt *et al*., 1996), CVlog in (Ruigrok *et al*., 2011), or the coefficient of local variation, Lv, in (Shinomoto *et al*., 2005). However, we see the quantities FF and CV^2^ as the best variability measures, and in this paper we present alternative estimators of their common value. These estimators can be considered as estimators of the local value of FF (local or instantaneous variability). Thus, we are not creating new measures of local variability, but new estimators of the local value of FF (CV). Note that an instantaneous version of CV is sometimes created by calculating of CV in the standard way, but using only a fixed number of ISIs around a given time (Borges *et al*., 2023; Prut & Perlmutter, 2003). However, such a method may have problem with bias due to small amount of data, as described above, or to be insufficiently local. We therefore propose a different approach.

We assume that the data used to estimate the variability are parallel spike trains (see Fig. 1). Such data are common in neuroscience and are usually obtained from repeated experimental trials or from simultaneous multi-unit neuronal recordings. However, recording all spike trains arriving at a target neuron is still not technically feasible. We are mainly interested in the short-term variability estimated based on this type of data. Our approach is such that we want to estimate variability based only on ISIs that include a particular time *t*_0_ that we shall construct (see Fig. 1). In the case where *t*_0_ is selected independently of the times of spikes (randomly), the ISIs containing the time *t*_0_ have in general different probability distribution than the ISIs observed from the time of their beginning. The difference between the distributions can be used to estimate FF. Based on this idea, we have created two variability estimators of FF. We derive their properties and show that they are surprisingly accurate and very easy to use. One of them is problematic when the probability of occurrence of short ISIs is high. This probability can be partially reduced by assuming a refractory period, and is not a problem at all in the second estimator. They are especially effective for estimation of instantaneous variability, where they significantly outperform the standard estimators.

**Figure 1.**
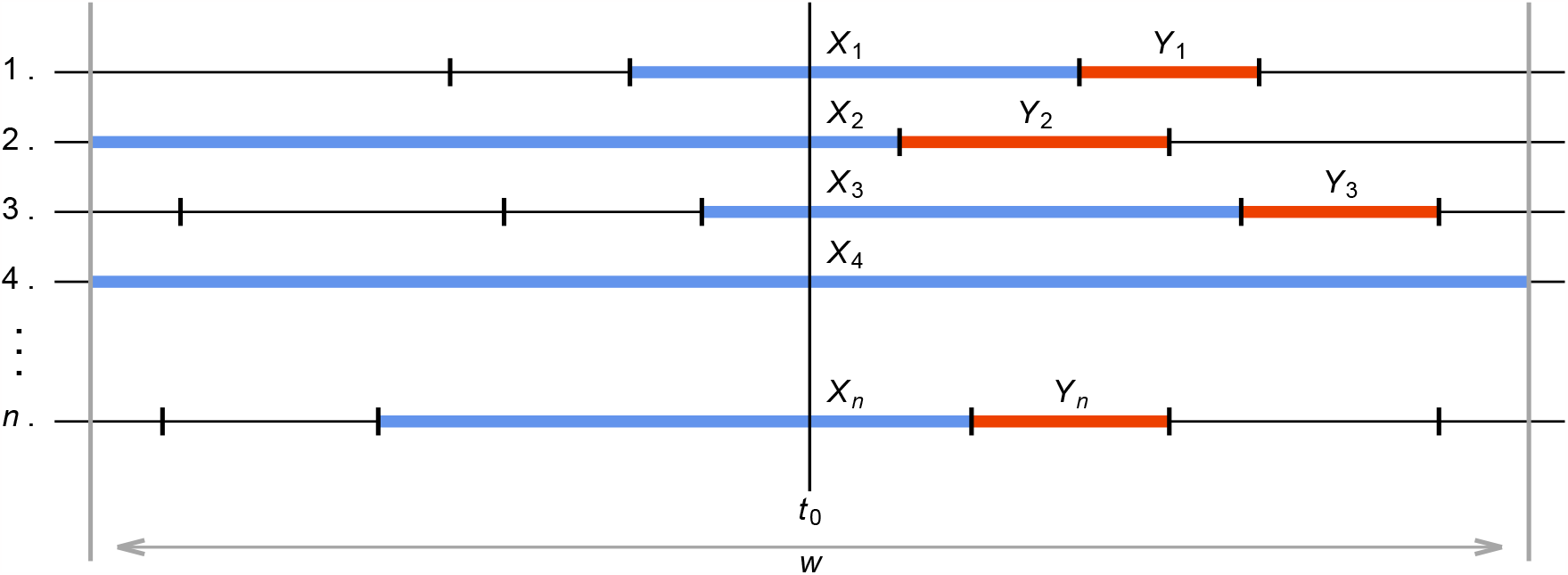
Illustration of the situation investigated in this paper. Assume *n* independent trials (realizations) of a spike train described by the equilibrium renewal point process of interspike intervals (ISIs). The length of consequent ISIs is determined by the random variable *T* (not shown). Our aim is to estimate the variability at time *t*_0_ which is not generally related to spike times. We base our estimators on the *instantaneous* ISIs, described by random variable *X* with realizations denoted as *X*_*i*_, that contain the time *t*_0_. In some cases, the immediately following ISI, *Y* with realizations *Y*_*i*_, is also employed. Note that due to the length-bias sampling the probability distribution of *X* generally differs from that of *T*, while the distributions of *Y* and *T* are the same. We exploit this fact and propose several novel estimators of the Fano factor (coefficient of variation of ISIs). In addition, the data used for the estimation may (but not necessarily) be restricted by the observation time window *w*.

The paper is structured as follows. Section 2 summarizes the necessary theory, defines the model of spike trains used (equilibrium renewal process), and the most important concepts and quantities - mainly the coefficient of variation, the Fano factor and their standard estimators. In Section 3, we propose the new estimators, given by equations (12) and (16), and derive their general properties. In Section 4, we consider several specific variants of the general spike train model (renewal processes with gamma, inverse Gaussian and lognormal distributions of the ISIs) and derive and illustrate corresponding properties of the new estimators. Maximum likelihood estimators of FF are also derived here. In Section 5, we use these models to compare the accuracy of the new estimators with the standard estimators using numerical simulations. In Section 6, we show a specific application - estimation of the change in the variability over time - and compare various estimators in this task. Finally, Section 7 summarizes the main results of the paper.

## 2. Spike trains model, variability measures and their estimators

We assume that the spike trains correspond to realizations of independent identical equilibrium renewal processes, i. e., the ISIs are identical and independent positive continuous random variables. The term equilibrium further implies that the time of the beginning of the spike trains does not affect the times of the current realizations of spikes, i.e., an infinite amount of time has passed since the beginning (Jewell, 1960). We denote the random variable describing the ISIs as *T* and its probability density function (pdf) as *f* (*t*). The mean and variance of *T* are denoted as *μ* = E(*T*) and Var(*T*). The intensity of the process is then (Cox & Lewis, 1966),

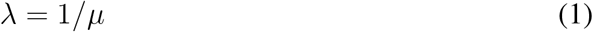

and the coefficient of variation is defined as

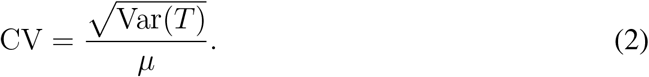

An alternative equivalent description of a renewal process is using the number of spikes up to a time *w >* 0, i.e., a counting process *N* (*w*). Based on this quantity, the Fano factor is defined as

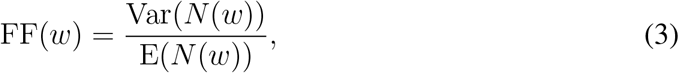

where time *w >* 0 can be seen as the length of an observation window. The term Fano factor is often used to refer directly to the limit

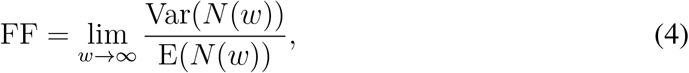

which removes the dependence on *w*. The Fano factor and the coefficient of variation are two of the most commonly used measures of the variability of neural spike trains (point processes). For renewal processes, they are related by a well-known equation (Cox, 1962),

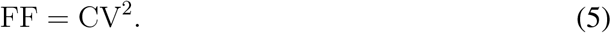

We also want to model and study the effect of the refractory period on the variability estimators. The (absolute) refractory period is a time interval of length *r* ≥ 0 that occurs after each spike, during which it is impossible to fire another spike. Such a property can be easily modeled by a renewal process by setting *f* (*t*) = 0 for *t* ∈ [0, *r*]. The refractory period is thus directly included in *T*, but sometimes it is useful to think of this random variable as

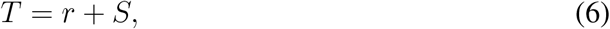

where *S* is the “random” part of *T*.

The sample counterparts of quantities (2) and (3) yield the standard estimators of the Fano factor and the squared coefficient of variation,

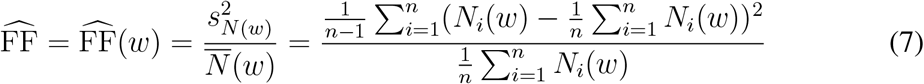

**Table 1:** Overview of the notation used in this paper. For the illustration see Fig. 1.

| Notation | Meaning |
| --- | --- |
| $T$ | Random variable representing lengths of ISIs (renewal process is used to model spike trains). |
| $f(t)$ | Probability density function of $T$. |
| $\mu$ | Mean of $T$, $\mu = E(T)$. |
| $\lambda$ | Intensity of the renewal process, $\lambda = 1/\mu$. |
| $w$ | Length of the observation window (used to get the spike counts), $w > 0$. |
| CV | Coefficient of variation. |
| FF($w$), FF | Fano factor, calculated in an observation window of length $w > 0$, and its limit for $w \rightarrow \infty$. |
| $r$ | Refractory period (dead time), $r \geq 0$. |
| $t_0$ | A time at which we want to estimate local variability. Time $t_0$ is generally not related to the spike (event) times. |
| $X$ | Random variable representing the lengths of the ISIs observed at time $t_0$. Its probability density function is given by equation (9). |
| $n$ | Number of parallel spike trains, also number of ISIs used in the estimators at each time $t_0$. In general, the number of data samples. |
| $T_i, i = 1, \dots, n$ | Realizations of $T$. |
| $X_i, i = 1, \dots, n$ | ISIs observed at time $t_0$ (realizations of $X$). |
| $Y_i, i = 1, \dots, n$ | First complete ISIs after $t_0$ (equivalent to realizations of $T$). |

and

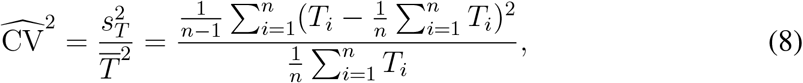

where *T*_*i*_ and *N*_*i*_(*w*) are the sample realizations of the random variables *T* and *N* (*w*) and *n* is the number of samples.

When observing the ISIs at (or around) a time *t*_0_ in *n* parallel spike trains, it is important to distinguish the relationship between the time *t*_0_ and the times of the spikes. One possibility is that *t*_0_ is completely unrelated to the actual spikes. Let us then denote the ISIs containing the time *t*_0_ as *X*_*i*_, *i* = 1, …, *n* (see Fig. 1). These ISIs generally do not correspond to the random variable *T* (since longer ISIs have a greater chance of being observed and vice versa), but to a random variable *X* with pdf (Kostal *et al*., 2018)

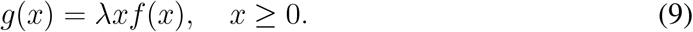

If we want to select ISIs corresponding to the random variable *T*, the selection of *t*_0_ cannot affect the length of the ISIs. E. g., we can select the first whole ISIs after *t*_0_. We denote such ISIs as *Y*_*i*_, *i* = 1, …, *n* (see Fig. 1). For better clarity of the used notation, there is a summary of the notation in Tab. 1.

## 3. Nonparametric (moment) estimators of instantaneous variability

Based on equation (9), we can derive

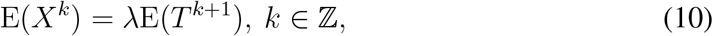

and

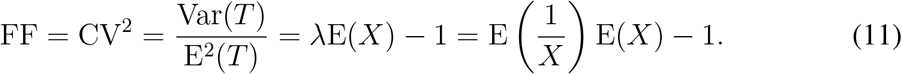

The value of FF can thus be calculated directly based on the mean of *X* and 1*/X*, yielding the estimator

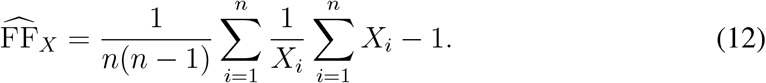

We interpret this formula as an estimator of FF based on *X*. The direct conversion of formula (11) to its sample counterpart would contain *n*^2^ instead of *n*(*n*− 1) in the first fraction. We made this change as it ensures that the estimator is then unbiased, i. e.,

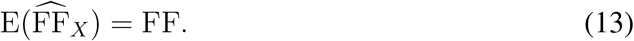

Note that both the standard variability estimators (7) and (8) have a bias depending on *n* that cannot be generally removed, they are only asymptotically unbiased (Rajdl & Lansky, 2014). Next, it can be derived,

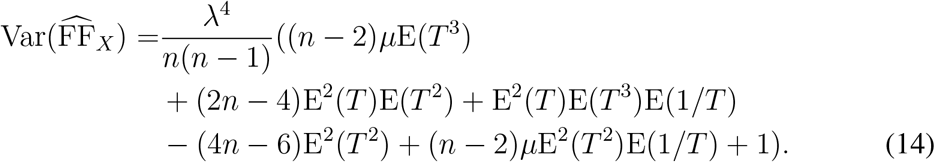

The variance (14) depends on the first few moments of *T* and can be thus calculated for a specific model of ISIs. The exact calculation of E(1*/T*) for *r >* 0 is usually problematic, but it can be approximated by numerical integration.

The mean of 1*/T* in formula (14) implies a potential problem of the estimator 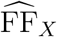, since it can diverge. E. g., E(1*/T*) is infinite even for the exponential distribution (Poisson process). In situations with large or diverging E(1*/T*), the obtained estimates are very unstable. In general, it holds that if *f* (0) *>* 0 then E(1*/T*) is infinite, and it can be proven that if *f* (*t*) is continuous on (0, ∞) with *f* (0) = 0 and *f* (0) exists and is finite, then E(1*/T*) *<* ∞ (Piegorsch & Casella, 1985). While the proof does not provide the complete sufficient and necessary conditions for the existence does not give complete conditions for the existence of E(1*/T*), but it confirms the intuitive idea that models with a non-negligible probability of occurrence of very small ISIs are problematic. Thus, we can expect that adding a refractory period to a renewal model can reduce the value of variance (14) (especially when the value of E(1*/T*) is high or infinite). We will explore the effect of the refractory period in specific illustrations later.

Moreover, we can eliminate this problem entirely by using the fact that E(1*/X*) = *λ* = E(*N* (*w*))*/w* and modifying formula (11) to

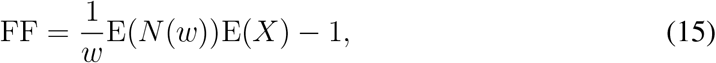

which yields a different estimator

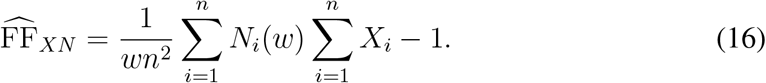

This estimator does not require the calculation of E(1*/X*), but it is not based purely on information obtained from *X*, it also uses information from the counts of spikes in a time window of length *w*. The window *w* must be somehow specified, a reasonable choice is to use a window that contains on average the same amount of “time” as used by 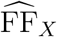, i.e.,

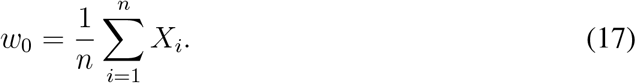

Estimator (16) then transforms into an interesting formula

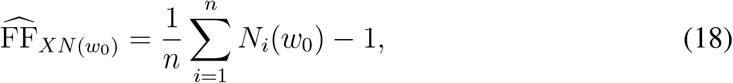

showing that FF can be simply estimated as the mean number of spikes in window of a length equal to the mean of the ISIs *X*_*i*_, 1 = 1 …, *n*, minus one. Note that formula (16) is also based only on the mean of two simple quantities although it is an estimator of variability.

Next we can derive

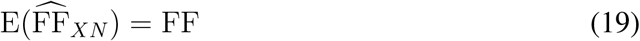

and

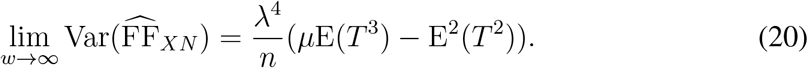

Differentiating equation (20) by *r*, assuming equation (6), yields (using Jensen’s inequality)

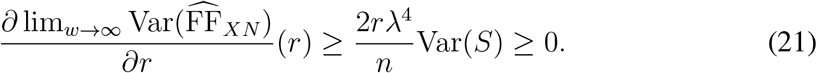

The variance of 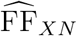 (for *w* →∞) is thus an increasing function of *r* and the minimum occurs for *r* = 0.

## 4. Probabilistic ISI models and MLE of instantaneous variability

For specific illustrations we use three probability distributions of ISIs (*T*) - gamma (GM), inverse Gaussian (IG) and log-normal (LN), including also the refractory period. They are probably the most often used distributions for modeling ISIs (Ditlevsen & Lansky, 2011; Nawrot, 2010; Shimokawa *et al*., 2010). All of these distributions are parametrized using two independent parameters (except for *r*), which can be expressed using *λ* and FF. The density of the gamma distribution is

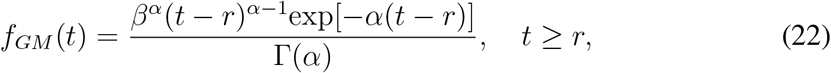

where *α* = (1 − *λr*)^2^*/*FF and *β* = *λ*(1 − *λr*)*/*FF, the density of the IG distribution is

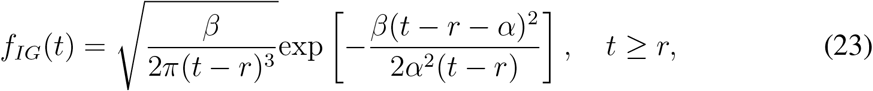

where *α* = 1*/λ* − *r* and *β* = (1 − *λr*)^3^*/*(*λ*FF) and the density of the LN distribution is

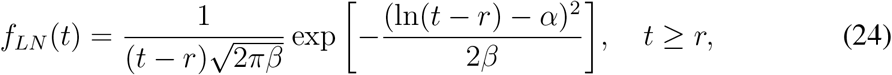

where *β* = ln(FF*/*(1 − *λr*)^2^ + 1) and *α* = ln(FF)*/*2 − ln(*λ*^2^(exp(*β*) − 1))*/*2 − *β/*2. For all the probability distributions it holds *r* ≤ 1*/λ* and *f* (*t*) = 0 for *t < r*.

Explicit formulas for 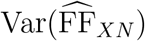 can be derived for *r* = 0. For the gamma distribution, equation (14) is then simplified to

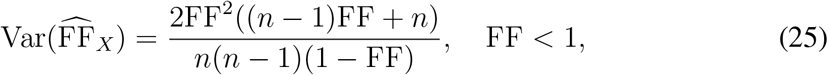

for the IG distribution to

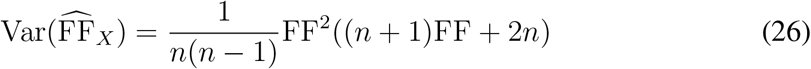

and for the LN distribution to

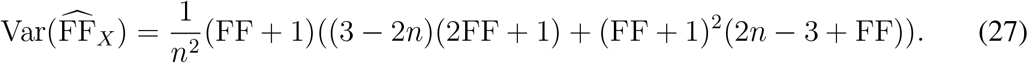

For the variance of FF_*XN*_, equation (20) gives the following relations. For the gamma distribution

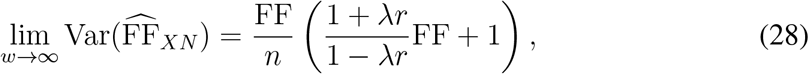

for the IG distribution

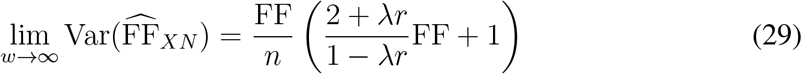

and for the LN distribution

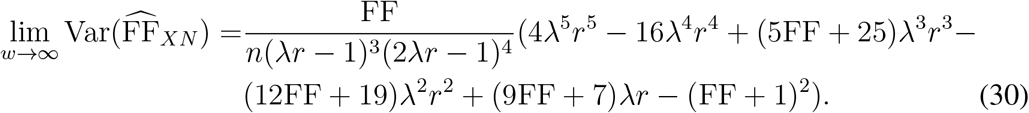

Note that formulas (28), (29), and (30) hold only for *w*→ ∞, but since *w* only affects the intensity estimator in formula (16), these approximations should give a good insight into the behavior of 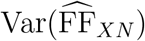.

Fig. 2 and 3 show the values of error of the estimators for the GM, IG and LN distributions of the ISIs as a function of FF and *r*. They are calculated based on equations (14) and (20), using numerical integration of E(1*/T*) if necessary. Specifically, we use the relative root mean square error,

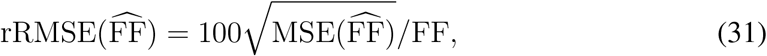

where 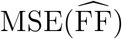 (mean square error) can be calculated as

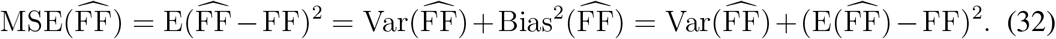

**Figure 2.**
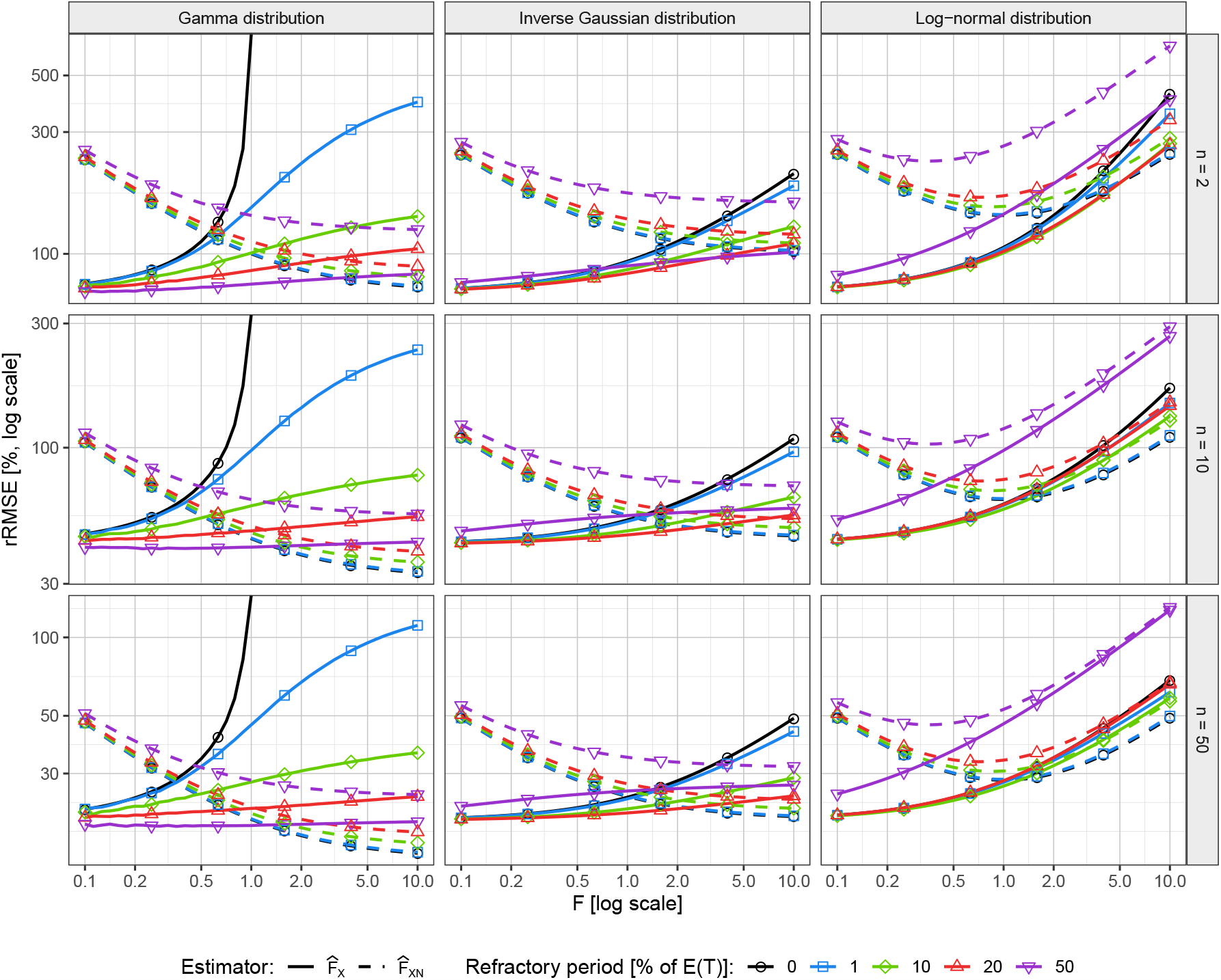
Performance of the proposed Fano factor estimators, 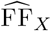 (solid line) and 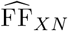 (dashed), measured by the relative root mean square error, rRMSE, in percent. The estimation is tested on three standard renewal spike train models (columns) given by the gamma, inverse Gaussian and log-normal distributions, in dependence on the number *n* of trials (rows) and on the true value of the Fano factor (horizontal axis). The color distinguishes the relative value of the refractory period (dead time) with respect to the mean ISI E(*T*). For 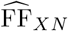 the values of rRMSE were calculated using the “ideal” infinite observation window. See Section 4. for the discussion of estimator performance.

**Figure 3.**
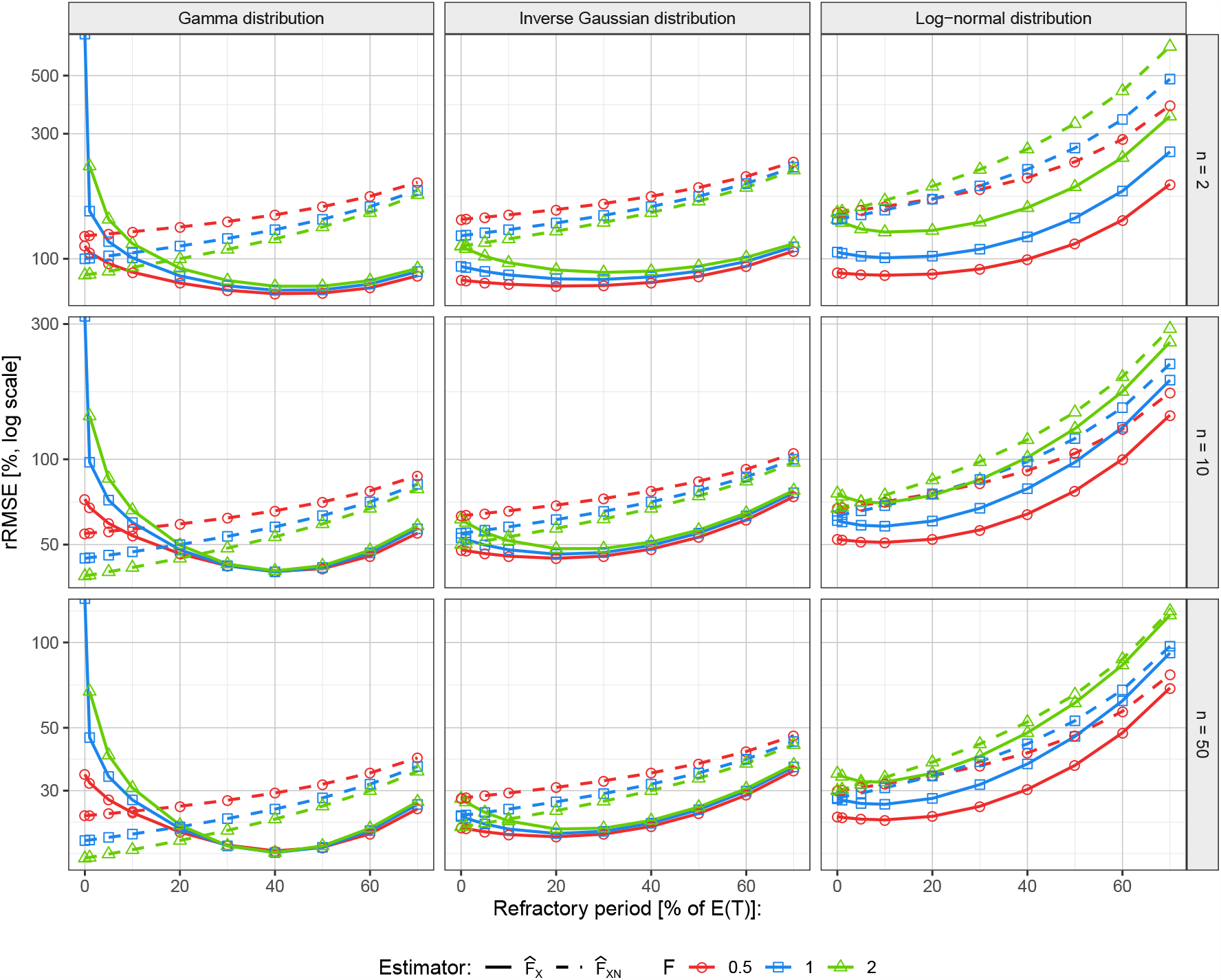
Performance of the proposed Fano factor estimators 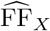 and 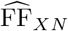. Visualization is analogous to Fig. 2. The estimation is tested for three different values of the true Fano factor (color) in dependence on the relative value of the refractory period (horizontal axis).

Let us look mainly at the dependence of rRMSE on the refractory period. For 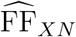, increasing the refractory period increases the error, in accordance with equation (21). For 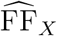, large values of *r* also have a negative effect on rRMSE, but smaller values reduce the error. This corresponds to the earlier described expected decrease of the value E(1*/T*) in equation (14). Thus, there is a minimum of the error for an *r >* 0. Specifically, for the gamma distribution with *r* = 0 the error is infinite for FF ≥ 1, since E(1*/T*) diverges, but this problem can be removed by adding a refractory period. E. g., with a refractory period of length 10% of *μ* the errors are reasonably small. In general, it seems that 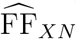 is more accurate for low values of FF and 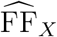 is more accurate for high values of FF, neither of which completely outperforms the other. With a low probability of very short ISIs, 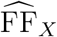 is often more precise. However, in the opposite case, it can yield very high errors, 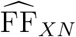 is then a safer choice.

Finally, we derive the maximum likelihood estimators (MLE) of FF based on realizations of *X* for the GM, IG and LN distributions of *T* (with *r* = 0). The derivation of the MLE assuming a gamma distribution of *T* leads to equation

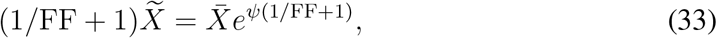

where *ψ*(*x*) is the digamma function, 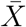 is the sample mean and 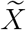is the sample geometric mean of *X*_1_, …, *X*_*n*_. Unfortunately, there is no analytical solution of equation (33), it must be solved numerically.

For the IG distribution, the MLE is directly equal to estimator (12), i.e., the moment and MLE estimators coincide. This is particularly interesting because the IG distribution arises as the first passage time of the Wiener process with drift, which is a widely used model of neuronal membrane potential. For the LN distribution, the MLE is

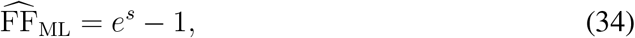

where

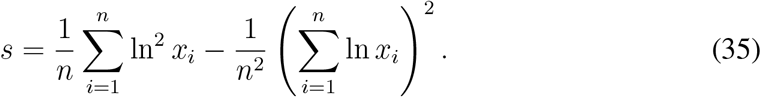

The above estimators were derived under the assumption that there is no refractory period (*r* = 0). For the case where *r >* 0, we present at least the MLE estimator for the exponential distribution. It is

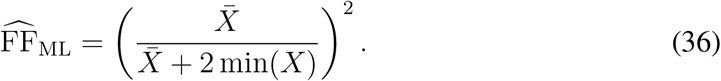

## 5. Comparison of 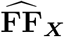 and 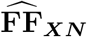 with the standard estimators

In the previous section, we have created two new estimators of FF based on the variable *X*. Now, we want to compare their accuracy with the standard estimators based on *N* (*w*) or *T*, given by equations (7) and (8). Let us now denote the estimator (7) as 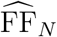 or 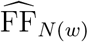 if we want to specify the used window. The question is then, what value of *w* should be used in this estimator, since the choice of *w* strongly influences its properties (in particular, smaller *w* yields mostly larger bias (Rajdl & Lansky, 2014)). We use mainly *w*_0_ given by equation (17). However, this window is selected based on realizations of *X*, so it may seem to favor the 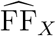 estimator. Therefore, we will also assume a (theoretically) infinite window, *w*_*I*_ = ∞ (very large in practice), which is the best choice for 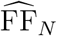. For estimator (8) we need realizations of *T*, so it is natural to use the first complete ISIs after *t*_0_ (values *Y*_*i*_, *i* = 1, …, *n*, see Fig. 1). We will denote this estimator 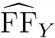 in the rest of the paper. In the estimator 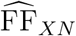 we use the window *w*_0_ to calculate the intensity. We calculate also the performance of the MLE estimators 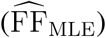, we always use the estimator derived for the specific distribution for *r* = 0, even if the real *r* is not zero.

To measure the accuracy, we again use rRMSE (31), but since there are no reasonably simple formulas for MSE for all the estimators, we will calculate the values using numerical simulations for spike trains with the GM, IG and LN probability distributions of the ISIs. For the parameters, we use the following values, *λ* = 1, *r* ∈{0, 0.1}, *n* ∈{2, 10, 50} and FF ∈ [0.1, 10]. For each combination of parameters, the values of error were calculated based on 20000 estimates for each estimator. The results for *r* = 0 are shown in Fig. 4 and for *r* = 0.1 in Fig. 5. Examining these figures yields the following conclusions.

**Figure 4.**
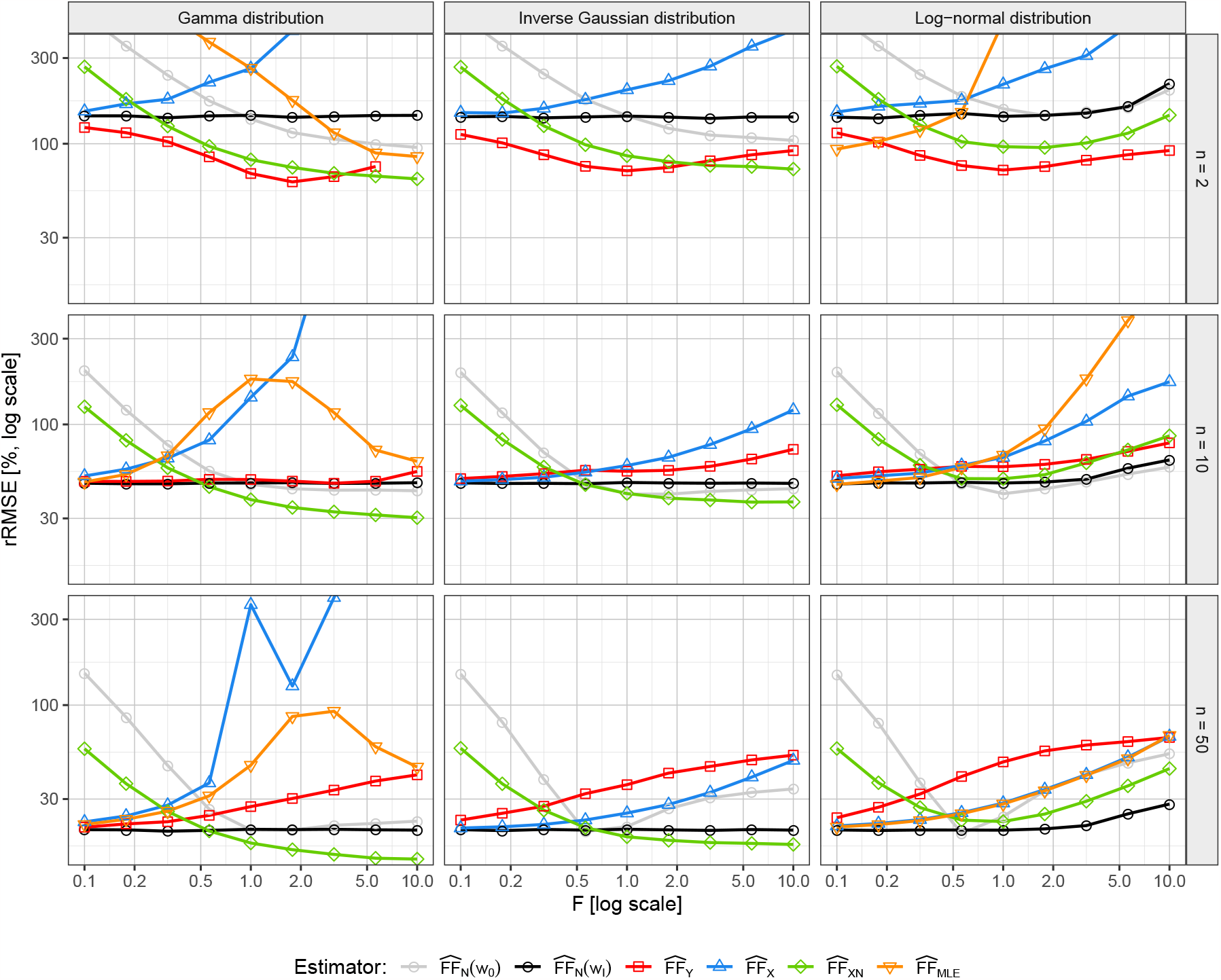
Comparison of performance between the proposed Fano factor estimators, 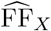 and 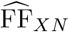, maximum-likelihood, 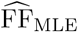, and non-parametric estimators based on standard formulas, 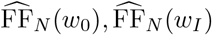 and 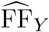. Analogously to Fig. 2 three standard ISI renewal models (columns) are tested at different number *n* of trials (rows) and various values of the true Fano factor (horizontal axis). For the purpose of the figure the refractory period is set *r* = 0, the mean ISI E(*T*) = 1 and each case was generated 20000 times. See Section 5. for more details and discussion of the results.

**Figure 5.**
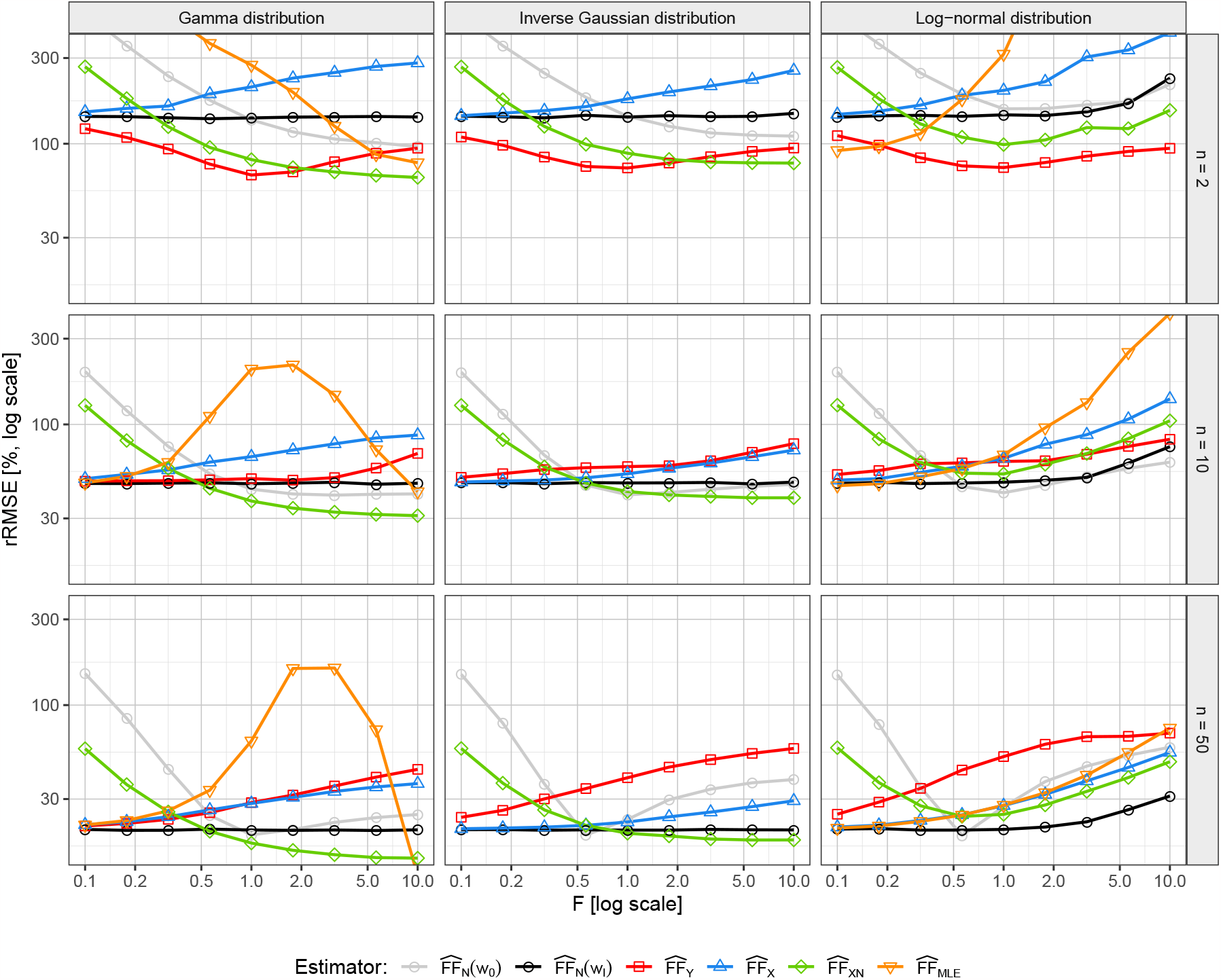
Comparison of various Fano factor estimators. Visualization and setting analogous to Fig. 4, except that the value of the refractory period is set to *r* = 0.1.

- RMSE of 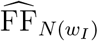 creates an almost constant line, representing approximately the lowest error that can theoretically be achieved by an 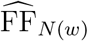 estimator. Surprisingly, it is rarely the best estimator.
- 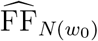 is relatively good (but not the best) for large FF, but mostly the worst for small FF.
- For very small *n*, 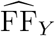 seems to be the best, but for larger *n* it is quickly outperformed by other estimators. In general, for the lowest *n* shown (*n* = 2), the errors are very large for all estimators.
- The error of 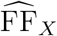 is often very small for small FF, but grows rather quickly with increasing FF. As expected, the refractory period reduces this error growth, especially for the gamma distribution.
- 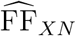 removes the problem of 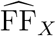for large FF, it is often the best for FF *>* 1, but the error increases for small FF. The refractory period does not affect the accuracy of this estimator too much.
- The MLE estimators for *r* = 0 have similar accuracy as 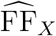. For the IG distribution the estimators are identical, for the LN distribution the errors seem to be very similar for large *n*, and for the GM distribution the MLE is better than 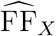 but also has problems with large FF.
- We also tested (not shown) the accuracy of the 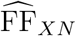 estimator while using the exact (not estimated) value of *λ*. Surprisingly, the resulting error is larger than when the intensity is estimated. The reason is that when both E(*X*) and *λ* are estimated, their errors (biases) can partially cancel each other out.
- The MLE (36) assuming an exponential distribution with a refractory period was also tested (not shown) for the situations presented. However, the results were poor, which is not surprising since the generated distributions are not exponential.

In conclusion, there is clearly no single estimator that is best in all situations, however, considering all the distributions and parameters, the 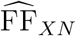 estimator seems to be the most accurate.

## 6. Estimation of the change in the variability over time

So far, we have assumed that we estimate the variability (Fano factor) at a specific (random) time from stationary spike trains. However, probably an even more important task is to estimate the time dependence of the Fano factor in multiple parallel spike trains to detect the changes in the (instantaneous) variability. The main advantage of the estimators 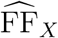 and 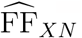 in this scenario is that there is no need to select a time window (except for the window for intensity estimation in 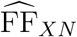), they use only one ISI from each spike train and thus automatically focus on the instantaneous variability. If we use *w*_0_ (17) as the length of the window for intensity in 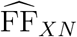, then even this estimator automatically adjusts the time scale it focuses on.

To test the accuracy of the estimators in such a situation, we need a suitable model of spike trains that allows us to change the variability (CV^2^, FF) at a specific time *t*_0_. Unfortunately, a renewal process cannot be directly adjusted to immediately change its variability at a given time. Therefore, we will generate two renewal processes, one with Fano factor FF_1_ and the other one with FF_2_, and join them at a random time *t*_0_. From the first process we will use the spikes up to *t*_0_, from the second process we will use the spikes after *t*_0_. Using this model, we test the ability of the studied estimators to correctly detect the change of FF over time. We assume *t*_0_ = 0, FF_1_ = 0.5 and FF_1_ = 2 and generate *n* = 50 parallel spike trains and estimate the Fano factor for a time sequence with a step of 0.05. We repeat the procedure 20000 times and compute the means and errors of the estimators. The estimators being compared are FF_*N*(1)_, FF_*N*(5)_, FF_*N*(10)_, FF_*Y*_, FF_*X*_ and FF_*XN*_. Three specific values of *w* (1, 5, 10) are used for the FF_*N*_ estimator to better see the influence of the size of the window. A comparison of these estimators for the GM, IG and LN distributions of the ISIs is shown in Fig. 6. We focus on the mean of the estimates, which should be as close as possible to FF_1_ for *t <* 0 and to FF_2_ for *t >* 0, and on rRMSE which of course should be as small as possible for all *t*.

**Figure 6.**
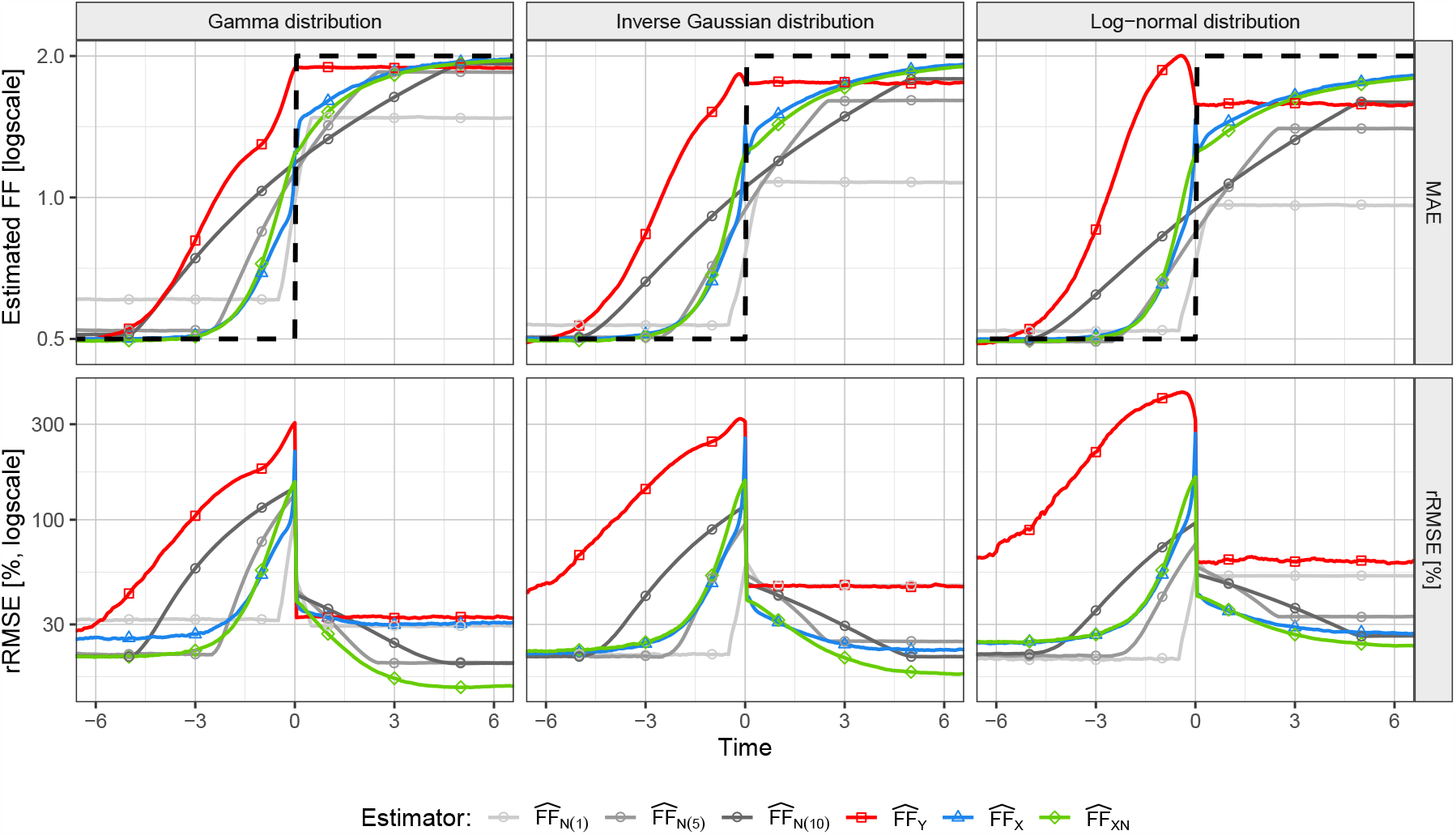
Estimation of spike train variability change in time. The proposed Fano factor estimators, 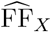 and 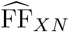, and standard non-parametric estimators, 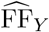 and 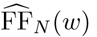 for three observation windows, *w*∈ *{*1, 5, 10}, are compared. The estimation is tested on three renewal spike train models with dead time (columns), the gamma, inverse Gaussian and log-normal distributions of ISIs, in dependence on time (horizontal axis). At time *t* = 0 the true Fano factor (black, dashed line) changes from FF = 0.5 to FF = 2. The top row shows the estimated Fano factor, the bottom row is the estimator performance measured by the relative root mean square error (rRMSE) in percent. The values were calculated from 20000 repetitions of 50 parallel spike trains with E(*T*) = 1 and refractory period *r* = 0.1.

Exploring Fig. 6 provides the following insights. The behavior of the 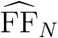 estimator is not surprising, for large *w* the estimated variability change is too smoothed, the estimator detects it too early. On the other hand, for small *w* the estimator has a large bias and there seems to be no satisfactory compromise. The 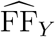 estimator is based on the first whole ISIs after the inspection time, so it catches the beginning of the change very poorly and generally has the largest error. It may seem strange that it has a large negative bias even for *t >* 0. This is a bias dependent on *n*, which vanishes for *n* → ∞, but is still large for *n* = 50. From the point of view of the mean values, the estimators 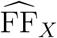 and 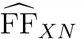 are very similar (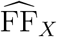 is slightly better) and quite good when compared to the other estimators. They capture the beginning of the variability change well and are the only estimators that converge to the true value of FF_2_ for *t* → ∞. From the point of view of rRMSE, 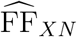 is mostly more precise than 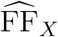 and even than the other estimators. For *t <* 0, 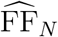 with smaller windows sometimes outperforms 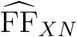, but for *t >* 0, 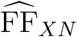 is clearly the best.

In summary, as in the previous section, no estimator is always the most accurate, but 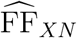 seems to be the best choice, especially for situations where we expect FF *>* 1.

The studied scenarios assume that the variability changes, but the intensity remains constant. However, in real experiments, the intensity is rarely constant, which could negatively affect the accuracy of the estimators. Therefore, we also explore situations where FF remains fixed, but the intensity changes. As the probability distribution of the ISIs, we use only the GM distribution, but with three values of FF (0.5, 1, 2). For *t*_0_ ≥ 0 the intensity is *λ*_1_ = 0.5, for *t*_0_ *>* 0 *λ*_2_ = 2. Other settings are analogous to the previous experiment. The results are shown in Fig. 7. Unfortunately, we can see that the new estimators are strongly influenced by the change in the intensity, which is mistaken for a change in variability. The standard estimators deal with this situation better, they are also influenced by the intensity change, but significantly less than the new estimators. One way to reduce this problem is to transform the spike trains into the operational time before estimating the variability. Operational time is such a time transformation that yields unit, constant intensity (time axis expressed in multiples of *μ*) (Nawrot, 2010; Rajdl *et al*., 2020).

**Figure 7.**
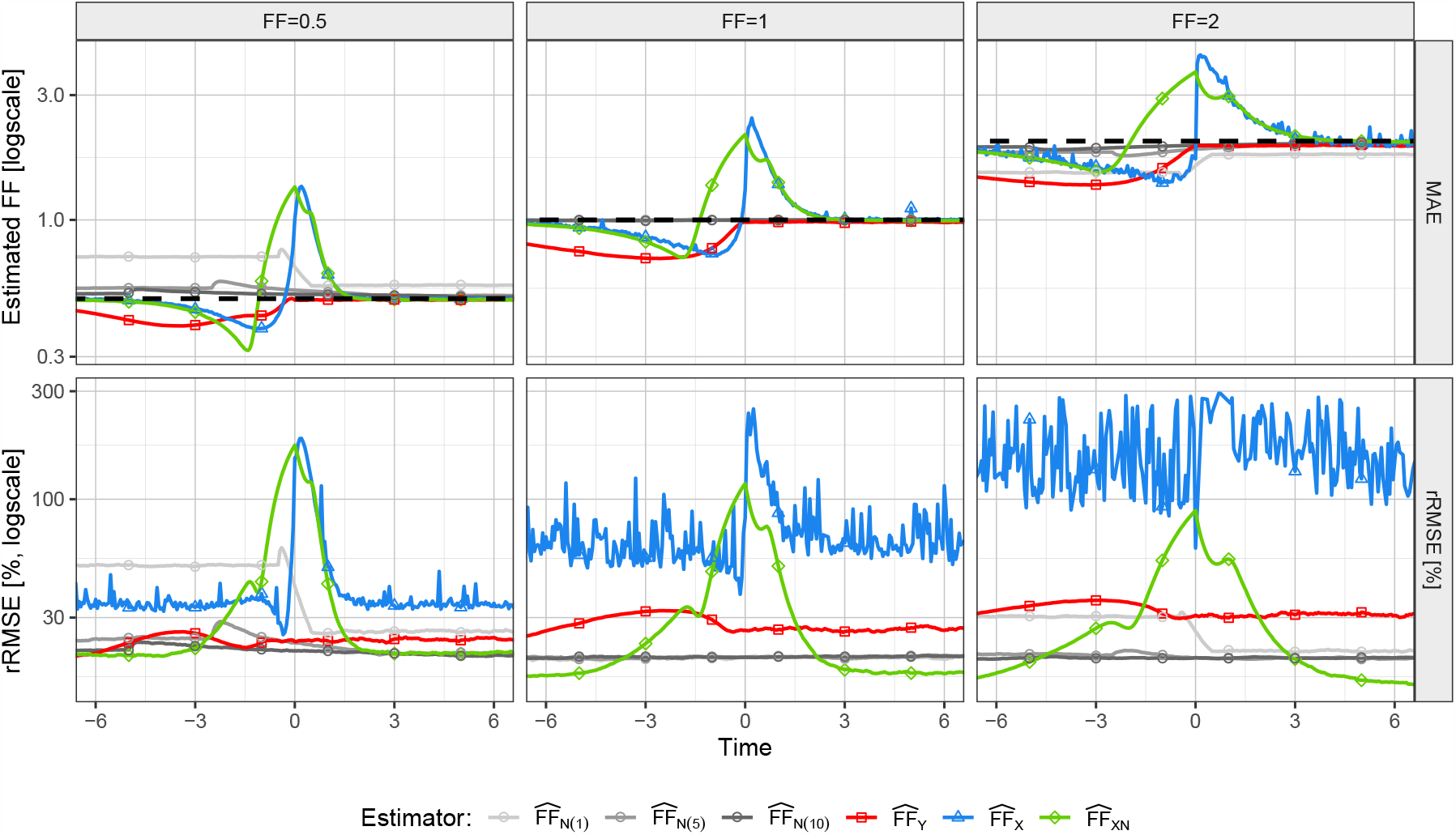
Estimation of spike train variability under change of firing rate in time. The proposed Fano factor estimators, 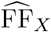 and 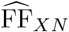, and standard non-parametric estimators, 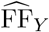 and 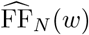 for three observation windows, *w* ∈*{*1, 5, 10}, are compared. The estimation is tested on the gamma distribution of ISIs (with dead time) such that there is a change in firing rate: E(*T*) = 2 up to time *t* = 0 and E(*T*) = 0.5 after *t* = 0. The true Fano factor however remains constant throughout each simulation (dashed line) at one of the three selected values (columns), FF = 0.5, 1, 2. The top row shows the estimated Fano factor, the bottom row is the estimator performance measured by the relative root mean square error (rRMSE) in percent. The values were calculated from 20000 repetitions of 50 parallel spike trains with refractory period *r* = 0.1.

## 7. Conclusions

We focused on the estimation of the (instantaneous) variability of multiple parallel spike trains at a specific, but random (with respect to the spikes) time *t*_0_. For such a task, standard estimators of the Fano factor or the coefficient of variation are often used. In the first case, the estimates are based on the number of spikes in a time interval around *t*_0_, in the latter case, the lengths of some ISIs are used (e. g., also from a time window). We showed that, under the assumption that the spike trains are realizations of a renewal process, it is very easy to estimate a quantity equivalent to the Fano factor based only on ISIs containing time *t*_0_ (one ISI from each spike train, see Fig. (1)), eventually to combine information from these ISIs with information from the number of spikes. In this way, we created two alternative Fano factor estimators 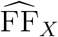 and 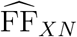 (which can also be seen as estimators of the coefficient of variation).

The first estimator, 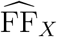, is based purely on ISIs *X*_*i*_, *i* = 1, …, *n*. Contrary to the estimators (7) and (8), 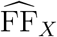 is unbiased and its accuracy is often better than the accuracy of the standard estimators. Another advantage is that when estimating the instantaneous variability, it is not necessary to adjust some parameters (e.g., the width of a window) to capture only local information. The estimator uses only one ISI from each spike train, thus automatically changes the used time scale based on the current intensity. It is also interesting that this estimator is the MLE estimator of FF assuming the IG distribution of the ISIs. The IG distribution can be derived as a distribution of ISIs from a very often used model of the neuronal potential – the Wiener process with drift, which gives another relevance to the 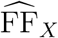 estimator. Unfortunately, it also has a relatively large drawback. Its error increases greatly for distributions of ISIs with a high probability of occurrence of very short ISIs. In the worst case scenario, the error can diverge, leading to very unstable estimates. We have shown that this problem decreases rapidly in the presence of a refractory period (which is common in real data), but it still cannot sufficiently improve every situation. We therefore modified the 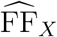 estimator to 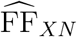, by replacing the problematic intensity estimation (part of 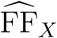 estimator) with an intensity calculated based on the number of spikes in a time window. It is then necessary to choose a time window for the intensity estimator, but we have proposed that the size of the window is simply calculated as the mean of the used ISIs. Such an estimator retains the good properties of 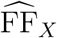, but completely removes the problem with short ISIs. Note also note that, although 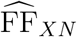 is a variability estimator, it is based only on the means of some simple quantities.

We derived some properties of the new estimators, mainly formulas for variance, and analyzed them. Then we performed various tests of the accuracy of the estimators, especially in comparison with the standard estimators (7) and (8). It turned out that no estimator is best in all situations, however, considering all the test scenarios, the most accurate is the new estimator 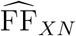. It is especially useful for estimating the time dependence of the variability. Besides being the most accurate, it is also very easy to use.

## Acknowledgements

This article is published with financial support from the Strategy AV 21 Programme, “Breakthrough Technologies for the Future – Sensing, Digitisation, Artificial Intelligence, and Quantum Technologies”.

